# Transient dopamine response on medium spiny neuron subtypes in switching approach-avoidance outcomes against action bias - A framework for exploration in action selection

**DOI:** 10.1101/2024.12.24.630270

**Authors:** Nitin Anisetty, Rohit Manchanda

## Abstract

An ensemble of direct and indirect pathway medium spiny neurons (dMSN and iMSN), compete via their neural activity to drive the decision to approach or avoid an object, respectively. Dopamine acting as a reward prediction error (RPE) signal causes experience-dependent synaptic changes in dMSN and iMSN, thereby shifting the dominance of neural activity to approach or avoidance signalling. These changes create bias in the striatal neuronal ensemble and restrict the choice to approach or avoidance in further iterations. However, organisms often exhibit behaviour where they choose undesirable or exploratory actions in anticipation of future reward or avoid desirable actions in anticipation of future risk. These against-bias decisions or exploratory decisions are not accounted for by the existing neuronal framework. To bridge this gap, we postulate that transient ‘motivational’ dopamine released from dopaminergic axons locally at sub-second timescales can cause temporary switch in dominance of neural activity between dMSN and iMSN leading to such adaptive decisions. By accounting for bias towards approach or avoidance or neither through synaptic weightages and accounting for differential affinity of DA to D1R and D2R, changes in dMSN and iMSN excitability at different levels of motivational dopamine was analysed. Furthermore, the spiking activity of striatal neuronal ensemble comprising of dMSN projecting directly and iMSN projecting indirectly onto the output nuclei of basal ganglia i.e. SNr neurons was simulated. This led to promising findings that demonstrate how SNr neuronal activity can generate outcomes that work against the cortico-striatal synaptic bias towards approach or avoidance.

**Graphical Abstract:** 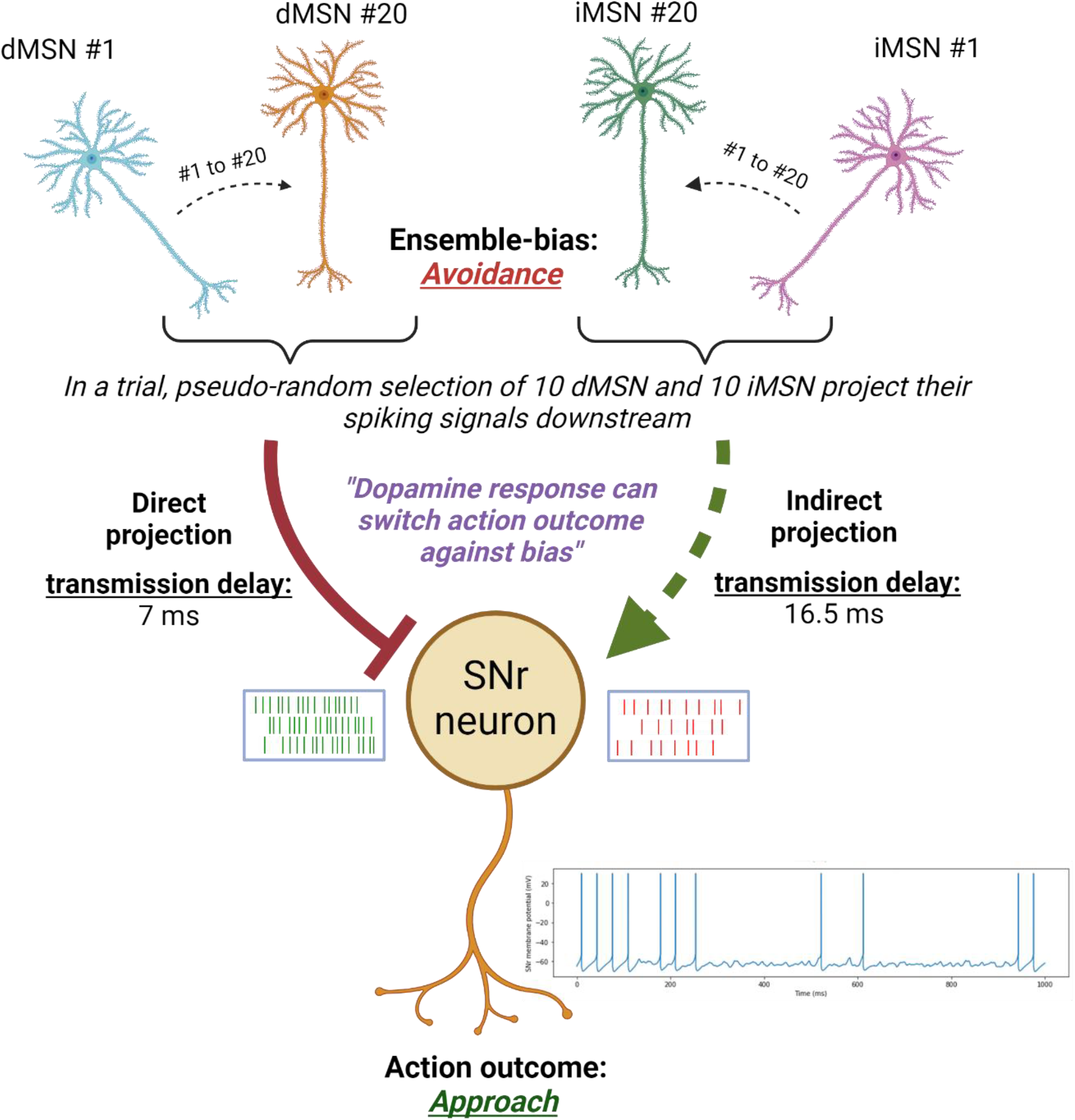

(Source: Created in BioRender. Anisetty, N. (2025) https://BioRender.com/a96f199)

## Introduction

To improve their chances of survival, organisms learn to differentiate between objects that are essential (such as food) and those that are dangerous (such as a predator). This allows us to repeat essential activities while avoiding dangerous ones. This widely researched process, referred to as value-based decision making, is driven primarily by the basal ganglia of the brain (Rangel et al., 2008; Orsini et al., 2019). In short, neuroplastic changes occurring especially in the input structure of basal ganglia (striatum) via the action of neuromodulator dopamine (DA) bias the neural activity to favour certain choices and avoid certain others (Meng et al., 2018; Bariselli et al., 2019; Cox and Witten, 2019). However, this schema fits only certain situations as organisms exhibit adaptability in decision making. Examples include avoiding high craving/value junk food to improve future health, or exploring a new option rather than exploiting known high craving/value options. Here, our main goal is to address this problem using a plausible mechanistic framework. To this end, we propose mechanisms of striatal neural activity leading to transient switch of decision, against the experience-derived bias, to approach or avoid an action.

The two main competing pathways in the basal ganglia, namely, the direct pathway and the indirect pathway, drive our decision whether to approach or avoid an action respectively, depending on which pathway dominates (Smith et al., 1998; Calabresi et al., 2014). A neural ensemble, comprised of ∼10 dMSN and ∼10 iMSN, encode and dictate whether an object should be approached or avoided (Barbera et al., 2016; Lee et al., 2020). As part of an action request, similar information pertaining to that action is received by a subset of dMSN and iMSN (Cui et al., 2013; Kress et al., 2013). According to the competitive model for action selection, when dMSN spiking activity dominates, thalamus is disinhibited causing approach behaviour. Likewise, when iMSN spiking activity dominates, inhibition of thalamus and default avoidance behaviour continues (Bariselli et al., 2019).

To account for adaptability in action selection, we consider as a possible explanation the role of *motivational dopamine (mDA)*. In early investigations, tonic firing of dopaminergic neurons was linked to motivation and was purported to take place on a timescale of seconds to minutes (Zhang et al., 2009). However, it was found recently that DA concentration can be altered in the nucleus accumbens at *sub-second timescales* as rats performed sequences of actions needed to achieve a reward (Mohebi et al., 2019). This mDA is released via local activation of dopaminergic axon terminals triggered by cholinergic interneurons and can be differentiated from the RPE DA that is released via somatic activation of dopaminergic neurons (Liu et al., 2022). Furthermore, cholinergic interneurons pause their firing during reward acquisition, thereby opening a temporal window for synaptic plasticity via RPE dopamine (Mohebi et al., 2023). The cellular mechanisms by which mDA exerts its effects are subjects of current conjecture and exploration. mDA could transiently act on dMSN and iMSN via several potential mechanisms including molecular cascades, genetic modifications and/or ion channels modulation. We hypothesize that to cause changes in the activity of MSN sub-types over sub-second timescales, ion channel modulation is the most plausible mechanism of action. Furthermore, transient DA modulation of ion channels in MSN has been reported (Nicola et al., 2000; Moyer et al., 2007) but its relevance to the larger scheme of action selection has not yet been elucidated. Collating these pieces of evidence, we hypothesize that mDA, acting as a transient signal, causes modulation of ion channels during selection of action, modulating dMSN and iMSN excitability either to reinforce an ongoing bias (for approach or avoidance), or to switch to an opposing signal, i.e. avoidance transiently dominates when the organism is biased to approach the object and vice versa. These changes in excitability of MSN sub-types can be attributed to how DA affects the state transitions and up-state dwell time (Anisetty and Manchanda, 2024).

Using the validated biophysical models of dMSN and iMSN from our recent work (Anisetty and Manchanda, 2024), we proceed to address questions of the kind that are best approached via this computational framework, viz, (i) comparing spiking frequencies of single dMSN v/s iMSN under different configurations accounting for biases (approach v/s avoidance) and differential affinities of dopamine to D1R v/s D2R, (ii) demonstrating switch in action outcome via SNr output neurons, under biased conditions, using ensemble activity of dMSN and iMSN. These questions are further adumbrated below.

The study can be broadly divided into two parts that are differentiated by the variation in the duration of synaptic input patterns. In the first part, the input frequencies of glutamatergic and GABAergic inputs switch every 284 ms leading to transition between states along this timeframe. This is to replicate the average up-state duration of 284 ms (Blackwell et al., 2003) and is also similar to preceding computational studies (Wolf et al., 2005; Steephen and Manchanda, 2009). Under this regime, spiking frequency was measured in dMSN and iMSN pairs that receive similar glutamatergic and GABAergic input patterns but differential dopaminergic input patterns due to varying affinities of DA for D1R and D2R. Furthermore, cortico-striatal plasticity is accounted for differentially in dMSN and iMSN, using different glutamatergic synaptic weights, to represent value learning via RPE response of DA. A key novel contribution here was that both models exhibited similar excitability under the tonic % DA receptor activation determined from literature.

In the second part of the study, the input frequencies of glutamatergic and GABAergic inputs switched once for 852 ms leading to a longer up-state along this timeframe. This long up-state reasonably represents the subject thinking of the action in question and whether to approach or avoid it. Within this duration we projected the spiking activity of the direct pathway and indirect pathway via dMSN and iMSN ensembles onto an SNr model neuron that indicates action selection in case there is a sufficient pause in its activity. Onset of pause and duration of pause were measured. The onset of pause could be crucial to determine which action is selected earlier among competing possibilities. Also, for a given configuration, we observe how different levels of DA % receptor activation can lead to change in action onset or duration of pause.

## Methods

As the premise of this study involves the interaction of transient dopamine with several ion channels and synapses, a biophysically constrained models of dMSN and iMSN were used in order to capture this interaction at different locations in the cells. The morphological and biophysical details of the dMSN and iMSN models as well as the dopamine dynamics in these models were described in detail in the companion research paper (Anisetty and Manchanda, 2024). Therefore, here we only focus on the novel aspects pertaining to this study.

### Relation between extracellular dopamine and % activation of dopamine receptors

To test that motivational dopamine (mDA) can cause opposing signalling, certain factors needed to be accounted for a more physiologically relevant interaction between DA and the MSN cell-types (dMSN and iMSN). Firstly, it is to be acknowledged that 0% DA receptor activation (either D1R or D2R) is not a baseline control condition as physiologically there is tonic activity of dopaminergic neurons leading to a non-zero baseline DA concentration and hence a non-zero baseline activation of dopamine receptors. This will be referred as the baseline % DA activation.

Secondly, due to the anatomical proximity of dMSN and iMSN that are encoding and competing for the same action, the dMSN and iMSN are also exposed to a similar extracellular concentration of dopamine (DA) (Klaus et al., 2017; Lee et al., 2020). Once DA is released into the extracellular space, there are several factors that affect the binding of DA to its receptors in the vicinity. DA mediates volume transmission, i.e., it spreads non-specifically in the extracellular space. The affinity of D2 receptor (D2R) is reported to be stronger than the affinity of D1 receptor (D1R) for DA (Rice and Cragg, 2008) (Kress et al., 2013). Therefore, at tonic DA, % activation of D2R will be greater than that of D1R due to its stronger affinity for DA.

For our study, we are interested in knowing the outcome post consideration of these, and other factors, on the % activation of D1R and D2R on dMSN and iMSN, respectively. The % activation or occupancy of DA to D1R and D2R in a given volume were worked out in a computational study by considering factors such as affinity of D1R v/s D2R, volume transmission, release probability, DA reuptake, etc (Dreyer et al., 2010). It is important to note that the % activation in our model indicates the fraction of D1R or D2R receptors of the individual dMSN or iMSN that are activated. For example, 50% activation of D1R on dMSN at any given instant indicates that approximately 2508 D1R out of 5016 D1R on the cell are active. Considering these factors, in the framework of our study design, we have translated the increasing level of dopamine in the extracellular milieu to varying % activation of D1R and D2R.

As the concentration of extracellular DA increases from zero, DA binds more to D2R owing to its higher affinity followed by DA binding to D1R gradually. Therefore, based on reasons ascertained, 17 conditions (as illustrated in Fig. 1) were used to represent the differential % dopamine receptor activations in dMSN and iMSN. Although (Dreyer et al., 2010) has reported similar % activations for a few key conditions (highlighted boxes in Fig. 1), the rest have not been reported in literature. Therefore, using the reported conditions as reference, the rest of the values were extrapolated reasonably such that the tonic/baseline % DA receptor activation conditions were in the middle of the spectrum of conditions. Even though the exact % activations may vary in vivo, the pattern of dopamine binding to D2R followed by D1R was maintained. Furthermore, the sensitivity of excitability to small % changes in receptor activation was found to be low, thereby giving leeway to extrapolate such conditions without major errors quantitatively.

**Figure 1.**
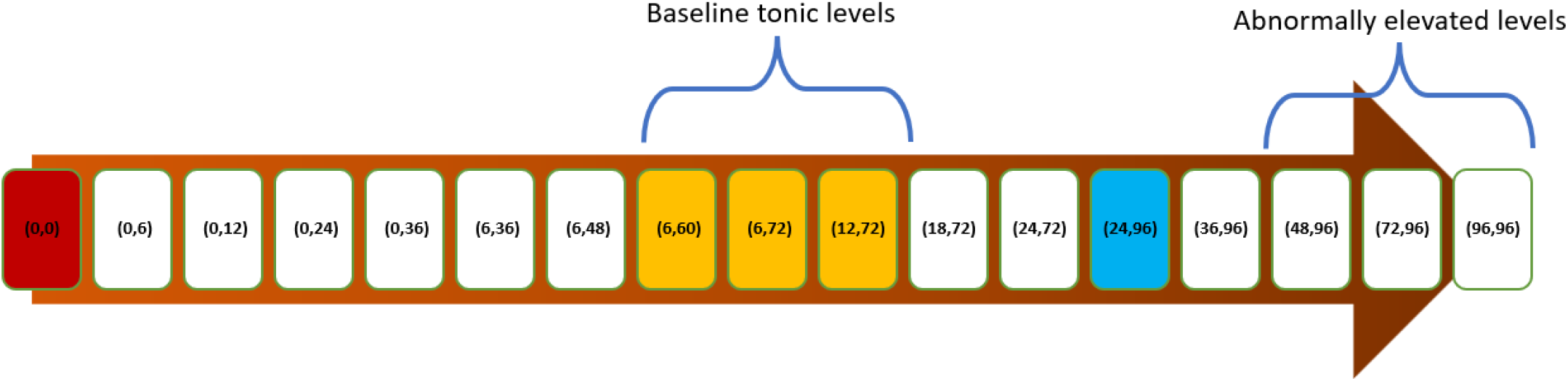
Increasing concentration of dopamine translating to different % activation of D1R and D2R. The pause, phasic firing, and tonic levels of % activation are shown in red, blue and yellow boxes respectively, similar to the reported values in (Dreyer et al., 2010) [Notation: (% D1R, % D2R)]

### Configurations to represent bias towards action and accounting for trial-by-trial variability

Randomization of synaptic input drive was introduced into the models to ensure trial-by-trial variability and to emulate physiologically realistic behaviour. However, to make the results replicable, we pseudo-randomize the simulations with the help of ‘seed’ variables. Each set of seed values will give a different sequence of randomization but the randomization sequence is replicable if we use the same set each time. The dMSN and iMSN models were subjected to glutamatergic, GABAergic, and dopaminergic synaptic inputs using asynchronous seeds for each trial. Three randomization seed values for input timings (of Glutamate, Dopamine, GABA inputs) and one randomization seed value for activated spine locations (to randomly select a subset of 5016 spines) were used for this study. For a given action, the inputs received on a single dMSN and a single iMSN in proximity are similar, hence the three timing seed values for both MSN sub-types were identical in each trial. However, the location of inputs and synaptic weights on dMSN and iMSN were always different.

In the framework of the competitive model, a dMSN and iMSN that are part of encoding a particular action were considered under 3 configurations as follows:

a. **No-bias configuration:** This configuration represents the idea of receiving an action request for a novel action. In this configuration, RPE dopamine dependent value-based learning has not occurred and the synapses of dMSN and iMSN have not been potentiated or depressed. Therefore, under tonic levels of dopamine, dMSN and iMSN should show similar excitability, indicating no tendency to approach or avoid the action.
b. **Avoidance-bias configuration:** This configuration represents the idea of receiving an action request that is biased for avoidance. This bias is due to prior exposure to the action and learning that the outcome of the action is distasteful. Therefore, under tonic levels of dopamine, iMSN has greater excitability than dMSN. Ideally, this action is to be avoided based on the organism’s experience but certain situations could warrant selection of this distasteful action.
c. **Approach-bias configuration:** This configuration represents the idea of receiving an action request that is biased for approach. This bias is due to prior exposure to the action and developing a propensity for it. Ideally, this action is to be selected based on the organism’s experience but certain situations could warrant avoidance of this high value choice.

The glutamatergic synaptic weights of dMSN and iMSN determine the bias towards approach or avoidance signal due to dopamine-dependent synaptic plasticity caused by previous experience with the encoded action. To incorporate this idea, a novel feature introduced in our models is variable glutamatergic synaptic weights, as opposed to constant value weights as used in previous models. Synaptic weights were increased by using random values picked from a certain distribution range to create no-bias, avoidance-bias, or approach-bias configurations of dMSN and iMSN based on distribution range used.

The model was highly sensitive to changes of weight values; hence we tried to restrict the values of synaptic weights from 1 to 1.1. For the no-bias configuration, the synaptic weights for dMSN and iMSN were picked from the same distribution range (1-1.05). In realistic situations, the synaptic weights of the two cell types will not be identical, therefore, even though the distribution range was the same for no-bias configuration, variability arises from using differing seed values. For the approach-bias configuration, dMSN synaptic weights were picked from a higher distribution range (1.05-1.1) and iMSN synaptic weights were picked from a non-overlapping lower distribution range (1-1.05). Similarly, for avoidance-bias configuration, iMSN synaptic weights picked from higher range (1.05-1.1) and dMSN synaptic weights were picked from lower range (1-1.05). ‘Higher synaptic weight’ (hsw) distribution indicates that the range of values is between 1.05 and 1.1 whereas ‘lower synaptic weight’ distribution indicates the range as being 1 to 1.05.

**Δfrequency (ΔF):** In order to draw comparisons between different % DA receptor activation conditions, we consider the use of a unified measure that streamlines the comparison. This variable is ΔF (= f_dMSN_ – f_iMSN_) which is the net difference in spiking frequency of dMSN and iMSN calculated for each trial. Therefore, for each condition, we get 10 values of Δfrequency corresponding to 10 paired trials of dMSN and iMSN.

### Izhikevich model for SNr neuron

Neurons of the Substantia Nigra pars Reticulata (SNr) are GABAergic and fire tonically with a frequency of ∼30 Hz (Sanderson et al., 1986; Gulley et al., 1999; Deransart et al., 2003; Maurice et al., 2003). To simulate its tonically spiking characteristics, the cell was modelled as an Izhikevich spiking neuron. The spiking threshold was taken as −51.8 mV to match with SNr neuron threshold as reported in literature (Richards et al., 1997). The neuron was also modified to accommodate voltage changes arising from excitatory and inhibitory post-synaptic currents (EPSCs and IPSCs). Inhibitory synaptic inputs arrived via the direct route from dMSNs and excitatory synaptic inputs arrived via the indirect route from iMSNs. Transmission delays via the direct pathway and indirect pathway were taken into account. For the indirect pathway route, total transmission delay was taken as 16.5 ms and for the direct pathway route it was taken as 7 ms (Lindahl et al., 2013).

**Table 1:** Parameters used for Izhikevich model of SNr neuron.

| Cell parameters |  | Post-synaptic current (PSC) parameters |  |  |
| --- | --- | --- | --- | --- |
| Parameter | Value | Parameters | EPSC | IPSC |
| *a* | 0.1 | $\tau_1$ (tau_1) | 0.65 | 0.5 |
| *b* | 0.2 | $\tau_2$ (tau_2) | 10.6 | 6.7 |
| *c* | -65 | $E_{rev}$ | 0 mV | -70 mV |
| *d* | 4 | weightage | 0.00145 | 0.0222 |
| *I* | 4.15 |  |  |  |

The IPSC and EPSC profiles of these inputs were validated based on data obtained via patch-clamp recordings on parasagittal brain slices of rats by (Bosch et al., 2011). WebPlotDigitizer was used to extract the experimental data traces. A two-state kinetic scheme synapse described by a rise time constant tau1 and decay time constant tau2 (derived from exp2syn of NEURON simulation environment) was used in the computational model. The parameters (time constant, weightage) for this were fine-tuned such that the simulated profile matched with the experimental ones (see Fig. 2).

**Figure 2.**
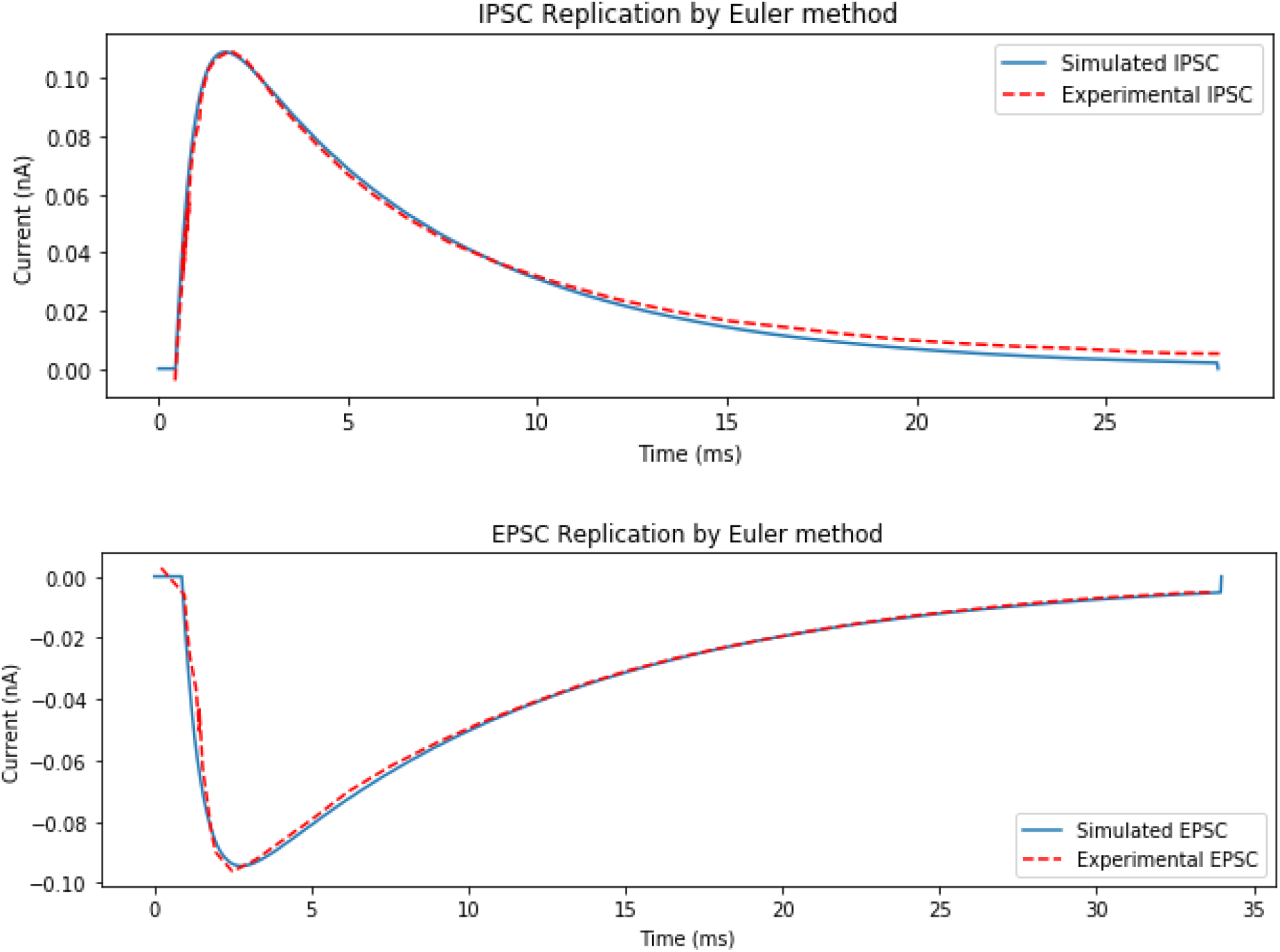
IPSC and EPSC profiles of SNr neuron validated with experimental data from (Bosch et al., 2011)

### Tracking ensemble response on SNr neuron

For a given action request, the ensemble of striatal projection neurons encoding the output response i.e. approach or avoidance are reported to be 10 dMSN and 10 iMSN on average (Barbera et al., 2016). To emulate this, we analyse the ensemble activity of 10 dMSN and 10 iMSN projecting onto SNr neuron. For each trial, these neurons are selected pseudo-randomly from a pool of 20 daMSN and 20 iMSN. In experiments, optogenetic activation of dMSN and iMSN leading to inhibition and enhancement, respectively, of the spiking activity of SNr neurons was shown by (Kravitz et al., 2010). However, it is to be noted here that only conduction delays across the indirect pathway have been accounted in our model giving way to further improvements in future studies.

To capture the dynamics arising during a choice to avoid or approach in a duration of 1000 ms, based on evidence that up-state switch occurs due to hippocampal inputs that encode episodic recollections, the duration of up-state was considered as 852 ms. This likely represents decision window and the up-state duration is within the physiologically range reported as 100-1000 ms (O’Donnell et al., 1999). Research indicates that the pause in SNr spiking activity during target selection in a saccade task varied in the range of 75-200 ms (Basso and Wurtz, 2002). Therefore, we considered the pause threshold in the mid-range i.e. 125 ms. The output measures used to quantify the output were the maximum duration of pause followed by onset time and duration of first pause if threshold was crossed.

## Results

### Effect of transient dopamine on spiking frequency of dMSN and iMSN

⁘ **Baseline % DA receptor activation of dMSN and iMSN:** As explained in methods, % receptor activation (D1R for dMSN and D2R for iMSN) at tonic levels of dopamine was considered as baseline. Furthermore, for no-bias configuration, we predict the spiking activity of dMSN and iMSN to be similar at these tonic levels. With no further fine tuning of parameters, we tested the dMSN and iMSN models to observe whether the speculated theory matches with simulated data. Using the Wilcoxon matched-pairs signed rank test, the spiking activity of dMSN and iMSN at (6,60), (6,72) and (12,72) conditions were found to be similar (n = 10, p>0.05) as seen in Fig. 3(a) and 3(b). This further strengthens the predictions of our models as the % tonic levels of DA receptor activation was arrived at from one strand of research (Dreyer et al., 2010) and the similarity in excitability at tonic DA under no-bias configuration lies at the foundation of a different strand of research (Hamid et al., 2016; Bariselli et al., 2019). Therefore, (6,72) condition was used as a standard condition to define configurations as approach-bias, avoidance-bias or no-bias. By using either higher or lower synaptic weight (hsw or lsw) distributions for glutamatergic synapses, the three configurations showed varied spiking activity at (6,72) condition.
⁘ **No-bias configuration (lsw-dMSN v/s lsw-iMSN):** As seen in Fig. 3(a), when levels of dopamine were lowered below tonic DA, reducing % receptor activation results in increased spiking frequency of iMSN while maintaining the spiking frequency of dMSN relatively similar as the binding of D1R does not change as much as that of D2R. On the contrary, increased spiking frequency for dMSN was observed while iMSN spiking frequency was maintained when levels of dopamine were raised above tonic DA, increasing % receptor activation (Fig. 3(b)). As the data did not pass the normality test in several cases, Wilcoxon matched-pairs signed rank test was conducted. The resulting spiking frequency data and p-values from the test were tabulated in Tables 2 and Table 3. Implications of the results are deliberated further in the discussion section.
⁘ **Approach-bias configuration (hsw-dMSN v/s lsw-iMSN):** Under this configuration, the spiking frequency of dMSN (11.8 ± 0.92) was significantly greater than that of iMSN (9.03 ± 0.74) at established baseline % receptor activation (6,72) indicating a tendency to approach. As seen in Fig. 3(c), with reducing dopamine levels leading to decreasing % receptor activation, the spiking frequency of iMSN first increases and becomes similar to dMSN followed by a switch in spiking activity in favour of iMSN having greater frequency than dMSN at very low dopamine levels. The spiking frequencies, especially at (0,36) and (0,24) are somewhat similar but above and below these conditions there is significant difference but in opposite direction. This data is tabulated in table 4.
⁘ **Avoidance-bias configuration (lsw-dMSN v/s hsw-iMSN):** Contrary to the approach-bias configuration, the spiking frequency of dMSN (8.38 ± 0.78) was significantly lower than that of iMSN (12.1 ± 0.85) at established baseline % receptor activation (6,72) indicating a tendency to avoid. As seen in Fig. 3(d), with elevated dopamine levels leading to increasing % receptor activation, the spiking frequency of dMSN first increases to become similar to iMSN followed by a switch in spiking activity in favour of dMSN having greater frequency than iMSN at very high dopamine levels. The spiking frequencies, especially at (24,72) and (24,96) are somewhat similar but above and below these conditions there is significant difference but in opposite direction. This data is tabulated in table 5.
⁘ **Δfrequency (dMSN – iMSN)**

The difference in spiking frequency between pairs of dMSN and iMSN that receive similar glutamatergic and GABAergic inputs were measured under different % activation conditions and under different bias configurations. As predicted, at baseline % DA receptor condition, the ΔF for no-bias configuration starts close to 0 whereas for approach-bias configuration it is higher. With decreasing % DA receptor activation, the pattern followed by no-bias and approach-bias configurations is decreasing. For no-bias configuration and approach-bias configuration, with decreasing % DA receptor activation, ΔF becomes more negative. The shift from positive ΔF to negative ΔF happens ∼ (0,36) % activation for approach-bias configuration whereas it starts off slightly below zero for no-bias configuration at baseline condition itself (Fig. 3(e)). On the other hand, with increasing % DA receptor activation for avoidance-bias configuration, ΔF shifts from negative to positive values eventually at ∼ (24, 96) % activation (Fig. 3(f)). The change in ΔF between every consecutive condition was statistically analysed using paired t-test with post-hoc Holm-Bonferroni correction (Table 6 and 7).

**Figure 3.**
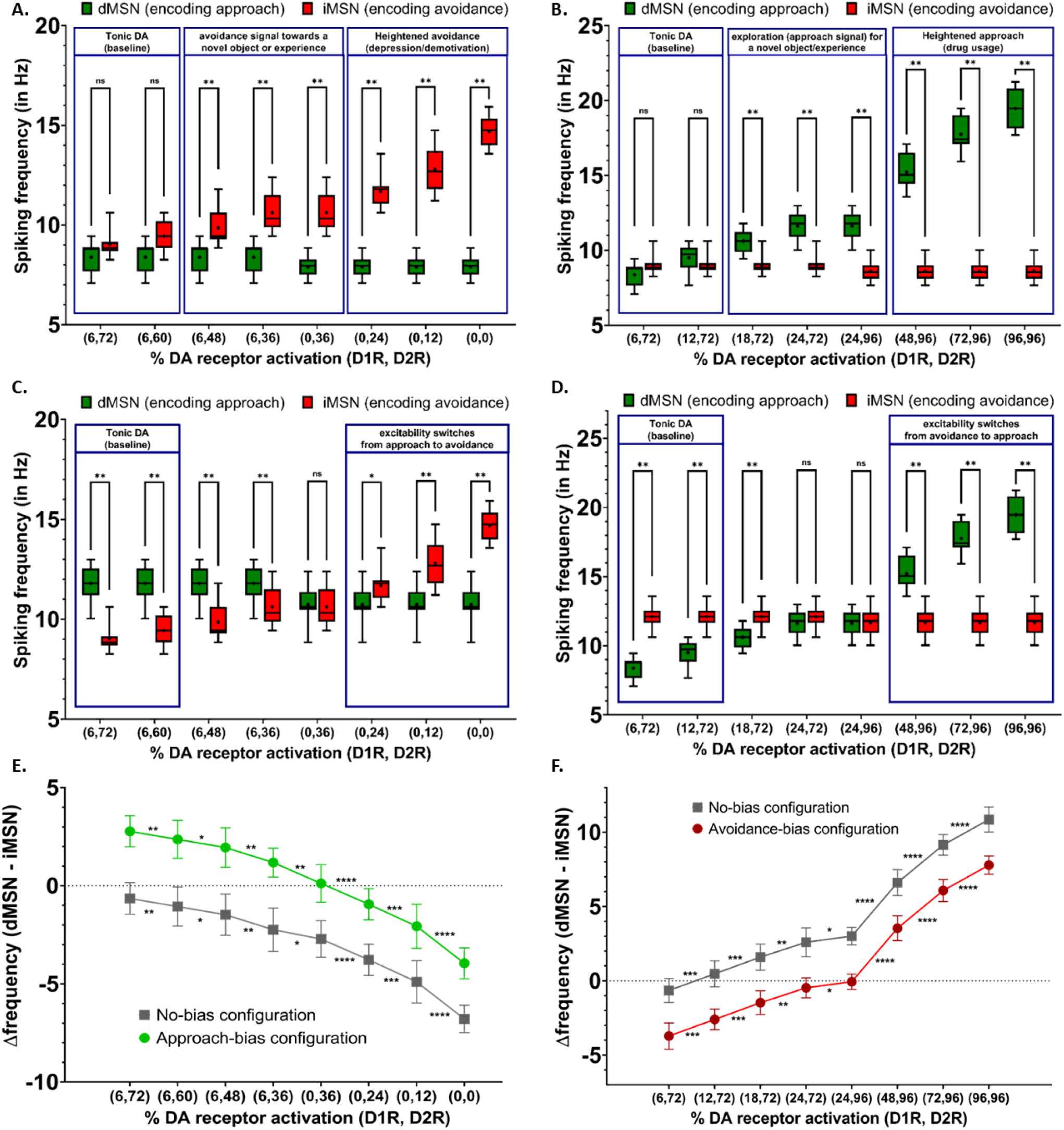
A) In the no-bias configuration, there is no significant difference in spiking frequency at tonic % DA activations but the avoidance signal starts to dominate as we decrease the % DA receptor activation along the x-axis. (B) In the no-bias configuration, as we increase the % DA receptor activation along the x-axis, we see that the approach signal starts to dominate. (C) In the approach-bias configuration, as we decrease the % DA receptor activation along the x-axis, we see that the approach-bias is overcome and the signal switches to avoidance. D) In the avoidance-bias configuration, as we increase the % DA receptor activation along the x-axis, we see that the avoidance-bias is overcome and the signal switches to approach. (E) Δfrequency for no-bias configuration and approach-bias configuration for decreasing % DA receptor activation (F) Δfrequency for no-bias configuration and avoidance-bias configuration for increasing % DA receptor activation

**Table 2:** Spiking frequency data of dMSN and iMSN in no-bias configuration under decreasing % receptor activation.

| % receptor activation<br>condition | Spiking frequency (mean ± SD) | | p-value |
| --- | --- | --- | --- |
|  | dMSN | iMSN |  |
| (6,48) | 8.38 ± 0.78 | 9.85 ± 0.93 | 0.0078 |
| (6,36) | 8.38 ± 0.78 | 10.62 ± 1.08 | 0.0039 |
| (0,36) | 7.91 ± 0.57 | 10.62 ± 1.08 | 0.002 |
| (0,24) | 7.91 ± 0.57 | 11.68 ± 0.87 | 0.002 |
| (0,12) | 7.91 ± 0.57 | 12.8 ± 1.11 | 0.002 |
| (0,0) | 7.91 ± 0.57 | 14.69 ± 0.81 | 0.002 |

**Table 3:**
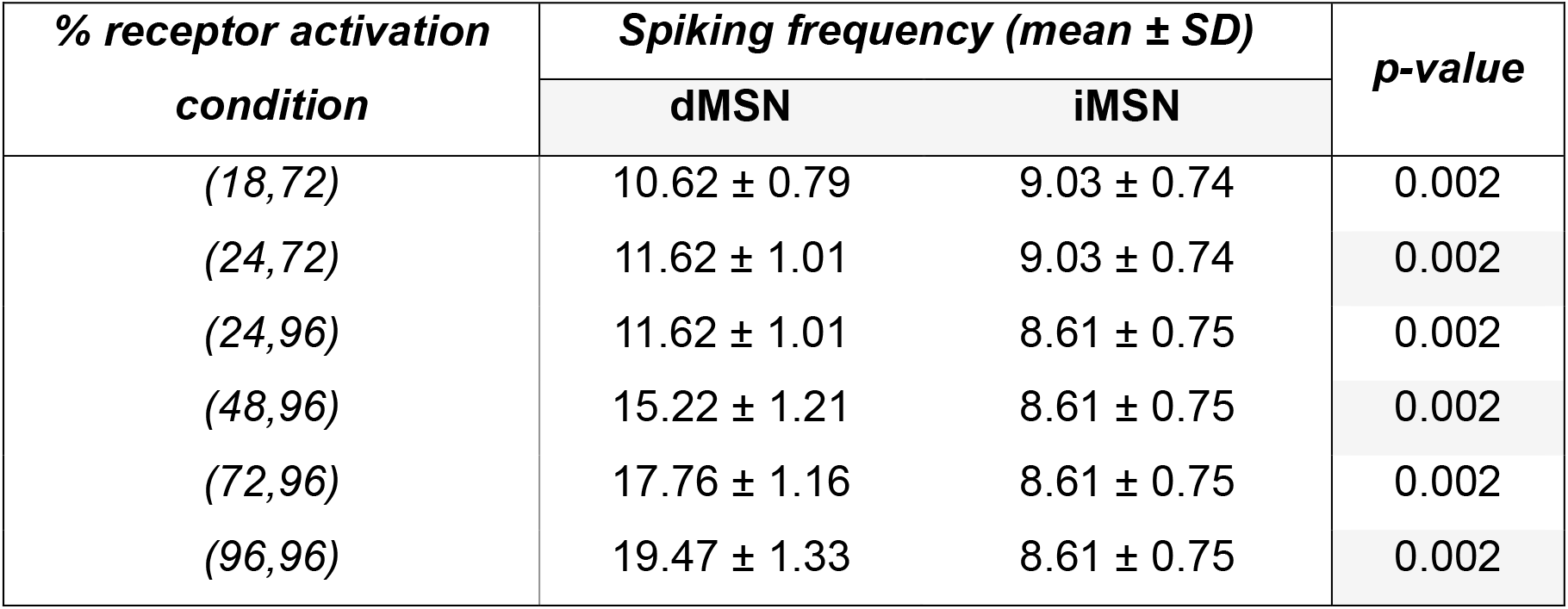
Spiking frequency data of dMSN and iMSN in no-bias configuration under increasing % receptor activation.

| % receptor activation<br>condition | Spiking frequency (mean ± SD) | | p-value |
| --- | --- | --- | --- |
|  | dMSN | iMSN |  |
| (18,72) | 10.62 ± 0.79 | 9.03 ± 0.74 | 0.002 |
| (24,72) | 11.62 ± 1.01 | 9.03 ± 0.74 | 0.002 |
| (24,96) | 11.62 ± 1.01 | 8.61 ± 0.75 | 0.002 |
| (48,96) | 15.22 ± 1.21 | 8.61 ± 0.75 | 0.002 |
| (72,96) | 17.76 ± 1.16 | 8.61 ± 0.75 | 0.002 |
| (96,96) | 19.47 ± 1.33 | 8.61 ± 0.75 | 0.002 |

**Table 4:** Spiking frequency data of dMSN and iMSN in approach-bias configuration under decreasing % receptor activation.

| % receptor activation<br>condition | Spiking frequency (mean ± SD) | | p-value |
| --- | --- | --- | --- |
|  | dMSN | iMSN |  |
| (6,48) | 11.8 ± 0.92 | 9.85 ± 0.93 | 0.0078 |
| (6,36) | 11.8 ± 0.92 | 10.62 ± 1.08 | 0.0039 |
| (0,36) | 10.74 ± 0.96 | 10.62 ± 1.08 | 0.002 |
| (0,24) | 10.74 ± 0.96 | 11.68 ± 0.87 | 0.002 |
| (0,12) | 10.74 ± 0.96 | 12.8 ± 1.11 | 0.002 |
| (0,0) | 10.74 ± 0.96 | 14.69 ± 0.81 | 0.002 |

**Table 5:** Spiking frequency data of dMSN and iMSN in avoidance-bias configuration under increasing % receptor activation.

| % receptor activation<br>condition | Spiking frequency (mean ± SD) | | p-value |
| --- | --- | --- | --- |
|  | dMSN | iMSN |  |
| (18,72) | 10.62 ± 0.79 | 12.1 ± 0.85 | 0.002 |
| (24,72) | 11.62 ± 1.01 | 12.1 ± 0.85 | 0.0938 |
| (24,96) | 11.62 ± 1.01 | 11.68 ± 1.11 | >0.9999 |
| (48,96) | 15.22 ± 1.21 | 11.68 ± 1.11 | 0.002 |
| (72,96) | 17.76 ± 1.16 | 11.68 ± 1.11 | 0.002 |
| (96,96) | 19.47 ± 1.33 | 11.68 ± 1.11 | 0.002 |

**Table 6:** p-values for Δfrequency (ΔF) compared across consecutive decreasing % DA receptor activation.

| **Comparison** | **Approach-bias** |  | **No-bias** |  |
| --- | --- | --- | --- | --- |
|  | **p-values** | **adjusted $\alpha$ (Rank)** | **p-values** | **adjusted $\alpha$ (Rank)** |
| (6,72) (6,60) | 0.0095 | 0.025 (#6) | 0.0095 | 0.0167 (#5) |
| (6,60) (6,48) | 0.0248 | 0.05 (#7) | 0.0248 | 0.05 (#7) |
| (6,48) (6,36) | 0.0063 | 0.0167 (#5) | 0.0063 | 0.0125 (#4) |
| (6,36) (0,36) | 0.0028 | 0.0125 (#4) | 0.0107 | 0.025 (#6) |
| (0,36) (0,24) | <0.0001 | 0.0071 (#1) | <0.0001 | 0.0071 (#1) |
| (0,24) (0,12) | 0.0002 | 0.01 (#3) | 0.0002 | 0.01 (#3) |
| (0,12) (0,0) | <0.0001 | 0.0083 (#2) | <0.0001 | 0.0083 (#2) |

**Table 7:** p-values for Δfrequency (ΔF) compared across consecutive increasing % DA receptor.

| **Comparison** | **Avoidance-bias** |  | **No-bias** |  |
| --- | --- | --- | --- | --- |
|  | **p-values** | **adjusted $\alpha$ (Rank)** | **p-values** | **adjusted $\alpha$ (Rank)** |
| (6,72) (12,72) | 0.0002 | 0.0125 (#4) | 0.0002 | 0.0125 (#4) |
| (12,72) (18,72) | 0.0002 | 0.0167 (#5) | 0.0002 | 0.0167 (#5) |
| (18,72) (24,72) | 0.002 | 0.025 (#6) | 0.002 | 0.025 (#6) |
| (24,72) (24,96) | 0.0445 | 0.05 (#7) | 0.0248 | 0.05 (#7) |
| (24,96) (48,96) | <0.0001 | 0.0071 (#1) | <0.0001 | 0.0071 (#1) |
| (48,96) (72,96) | <0.0001 | 0.0083 (#2) | <0.0001 | 0.0083 (#2) |
| (72,96) (96,96) | <0.0001 | 0.01 (#3) | <0.0001 | 0.01 (#3) |

### Indirect effect of transient dopamine on SNr neurons via striatal ensemble projections

Although the previous section gives us strong evidence on how the excitability of dMSN and iMSN is affected by transient mDA, we further test whether this spiking activity can drive changes in the output nuclei of the Basal Ganglia, namely, the neurons of Substantia Nigra pars reticulata (SNr). In Fig. 4, the onset and max duration of pause are shown for trials where the action threshold of 125 ms was crossed. Max duration refers to the longest pause during that trial whereas onset of pause refers to first pause that is greater than 125 ms. In Fig. 4(A), approach-biased configuration was used for the striatal ensemble and reduction in DA % receptor activation led to avoidance outcome indicated by increased firing of SNr neurons. In Fig. 4(B), avoidance-biased configuration was used and increase in DA % receptor activation led to approach outcome indicated by significant pause in SNr activity.

**Figure 4.**
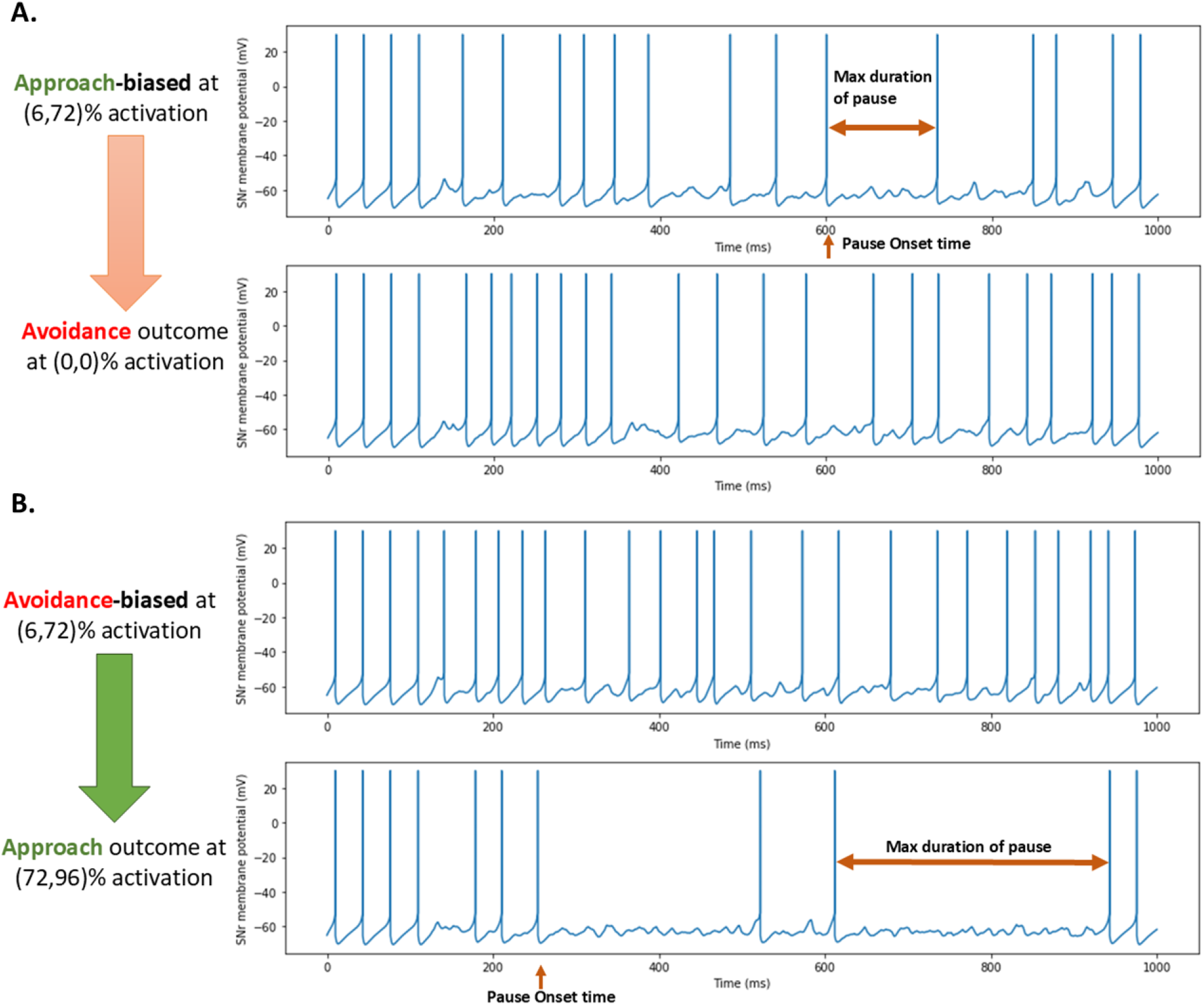
Switch in outcome against bias in SNr neuronal activity (A) Avoidance outcome observed with lack of significant pause in SNr activity at low dopamine levels in an approach-biased ensemble (B) Approach outcome observed with significant pauses in SNr activity at high dopamine levels in an avoidance-biased ensemble

The impact of spiking activity of dMSN and iMSN ensemble on SNr neurons across different % DA receptor activation conditions was analysed over 20 trials (Refer graphical abstract). The reduction of transient dopamine led to increase in SNr activity in all configurations which cannot be quantified as action threshold is not crossed. Therefore, such conditions were primarily considered only for the qualitative heat map of outcome (Fig. 6). In conditions where the threshold is crossed in majority of trials, the onset and duration of pause was analysed as shown in Fig. 5. Consecutive conditions were compared for significance using paired t-test with post-hoc Holm-Bonferroni correction (Table 8 and 9). The significance level was also indicated using symbols in Fig. 5(A) and 5(B).

**Table 8:** p-values for onset time of pause (Adjusted α following post-hoc Holm-Bonferroni correction indicated)

| *Comparison* | *Avoidance-bias* |  | *No-bias* |  |
| --- | --- | --- | --- | --- |
|  | **p-values** | **adjusted $\alpha$ (Rank)** | **p-values** | **adjusted $\alpha$ (Rank)** |
| (24,72) (24,96) | 0.0538 | 0.05 (#4) | 0.7896 | 0.025 (#3) |
| (24,96) (48,96) | 0.0162 | 0.025 (#3) | 0.0077 | 0.0167 (#2) |
| (48,96) (72,96) | <0.0001 | 0.0125 (#1) | 0.0008 | 0.0125 (#1) |
| (72,96) (96,96) | 0.0004 | 0.0167 (#2) | 0.9968 | 0.05 (#4) |

**Table 9:**
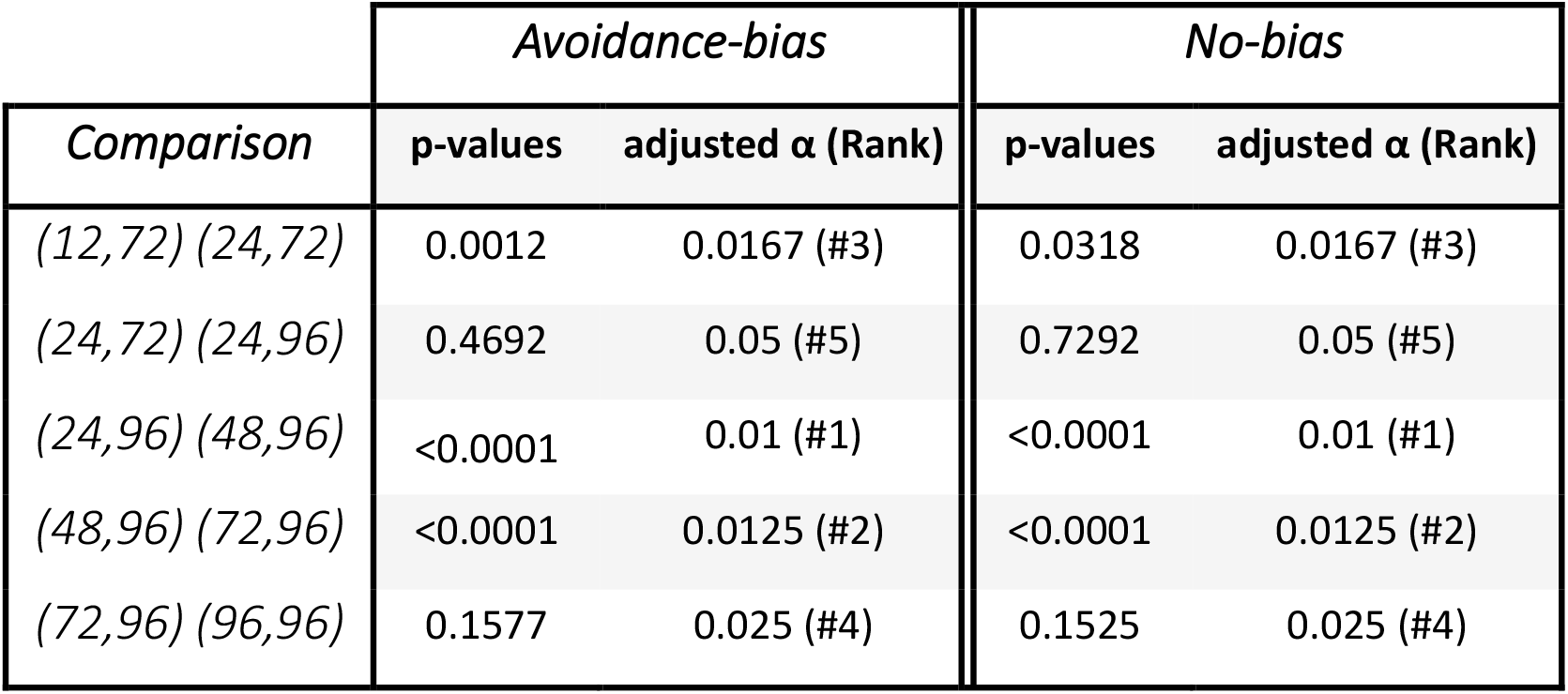
p-values for maximum duration of pause (Adjusted α following post-hoc Holm-Bonferroni correction indicated)

**Figure 5.**
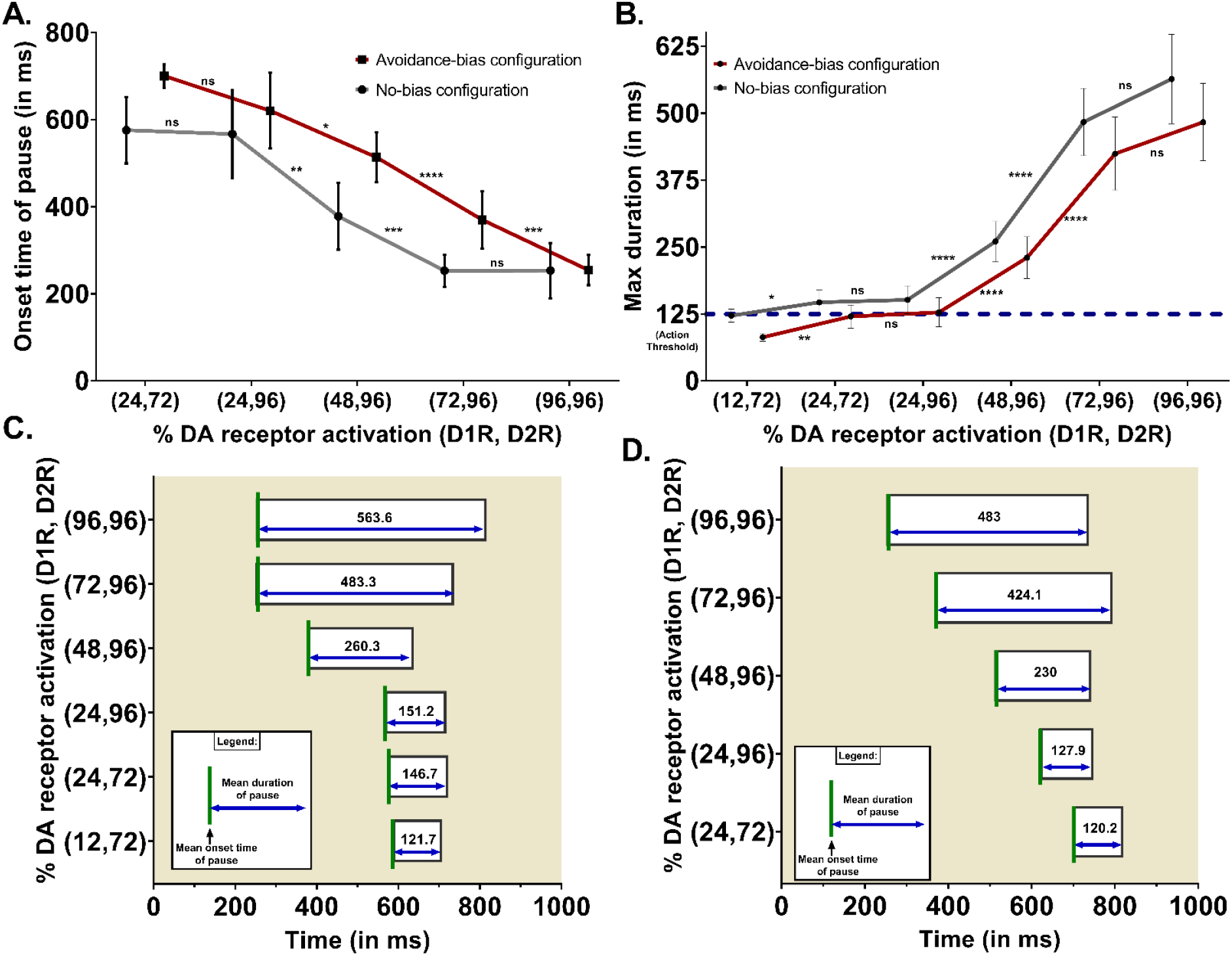
A) Mean with 95% CI was indicated [Note: When max duration of pause was below 125 ms there is no data for onset time of pause as we are referring to onset time only when threshold is crossed] Increasing DA leads to earlier onset and increased duration of pause in both (C) no-bias configuration and (D) avoidance-bias configuration

**Figure 6.**
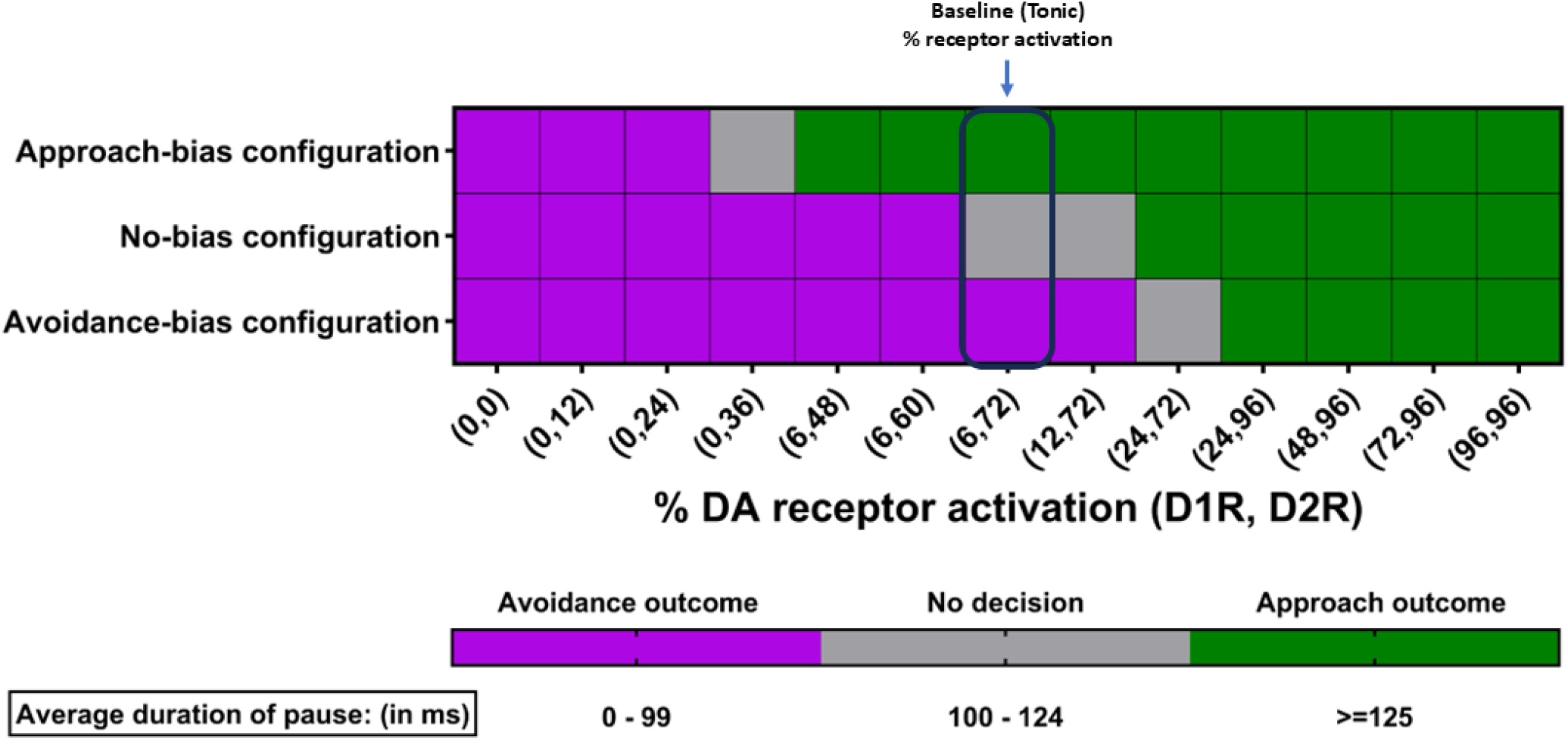
Qualitative heat map for maximum duration of pause averaged across several trials for each condition under three different bias configurations.

Earliest onset time observed is ∼253 ms on average and as expected it requires greater transient DA for avoidance-biased action (configuration) to reach this. The mean onset and mean duration of pause are visualized in Fig. 5(C) for no-bias configuration and in Fig. 5(D) for avoidance-bias configuration. With increasing activation of DA receptors, the mean onset time reduces and pause duration increases as predicted.

A qualitative heat map captures the decision outcome i.e. approach or avoidance or neither based on the maximum duration of pause averaged across 20 trials for each activation condition (Fig. 6). In cases where the pause threshold is not crossed, especially for lower DA receptor activation conditions, only few trials were used as results are qualitative. Interestingly, at tonic DA or baseline % DA receptor activation, all three configurations match with the outcomes they predict by definition. No decision zone represents a buffer window of neither decision occurring as <100 ms is definitely small for action being approached and >125 ms is a strong contender for action approach. From the qualitative heat map, we observe that action outcome pivots for approach-bias configuration at (0,36) % condition, for no-bias configuration at (12,72) % condition and for avoidance-bias configuration at (24,72) % configuration.

## Discussion

In our recent study (Anisetty and Manchanda, 2024), we developed biophysically constrained models of dMSN and iMSN to investigate the impact of transient dopamine on the state transition properties of these cells. The focus there was on the biophysically properties whereas for this study we expand our focus towards decoding exploration in action selection i.e. approach or avoidance. Biophysically detailed models of MSN sub-types were required for this study as the spatiotemporal patterns of dopaminergic receptor activations led to interwoven ion channel modulation dynamics that could not be captured otherwise. Additionally, these mechanisms are at the crux of our hypothesis that explains how actions can transiently switch to act against existing bias for approach or avoidance. In continuation to the previous work, we first analysed how the spiking frequency of MSN sub-types was affected in physiologically relevant conditions by accounting for factors such as differential DA receptor affinities and synaptic biases. Next, the spiking activity of several dMSN and iMSN, acting as a neural ensemble were projected directly or indirectly onto the output neurons of BG i.e. the SNr neurons. The key idea was to demonstrate that: a) the spiking activity of dMSN and iMSN can switch dominance against existing bias for approach or avoidance and b) this spiking activity of striatal ensemble can influence the outcome at the output neurons (SNr) to act against bias.

The striatum is known to have anatomical-functional correlation wherein, localized clusters are segregated as per their functioning for motor control or action selection (Alexander and Crutcher, 1990; Barbera et al., 2016). In this context, dMSN and iMSN that are anatomically proximal encode for the same function and are said to receive similar convergent synaptic inputs from other regions (Kress et al., 2013). Therefore, during any trial, for a given pairing of dMSN and iMSN, the glutamatergic and GABAergic input timings used were similar, however, the spatial location of activated dendritic spines were different. Although the extracellular milieu of dopamine shared by this dMSN-iMSN pair is same, the spatiotemporal % activation was different as per several factors such as differential affinity of DA for D1R and D2R, volume transmission of DA, etc on the basis of calculations by (Dreyer et al., 2010).

There is evidence indicating how a subset of dMSN and iMSN forming a neural ensemble project information regarding a certain action to determine the outcome as either approach or avoidance. Each such segregated and parallel processing streams were referred to as “channel” in the Basal Ganglia (Gurney et al., 2001; Romanelli et al., 2005). The synaptic changes induced on dMSN and/or iMSN by reward prediction error signals of dopamine biases the action channel towards either approach or avoidance (Gurney et al., 2015; Bariselli et al., 2019). This framework is incomplete as it does not explain behaviours that are exploratory in a broad sense. In this context, when there is no synaptic bias induced, the dMSN and iMSN spiking should be similar. Interestingly, under predicted tonic % activation of D1R and D2R, we observed balanced spiking in dMSN and iMSN. This is a crucial advancement in this study.

### Effect of transient dopamine on spiking frequency of dMSN and iMSN

In all three configurations, our findings indicate how changing levels of mDA, leading to differential activation of D1R and D2R, transiently alters the excitability of dMSN and iMSN that can either reinforce existing bias or potentially oppose the bias leading to approach signal instead of avoidance or vice versa. In case of no bias, transient mDA can influence the signal in either way depending on the direction of mDA change. Henceforth, we have referred to action specific information being sent via cortical and limbic inputs to the MSN sub-types as “action request”. The term request was used to indicate that the action is being considered but the outcome is not finalized at this stage.

#### No-bias configuration

In the no-bias paradigm, the action request received for processing, represents an unexplored action therefore there is no prior tendency for approaching or avoiding it. Note that it is possible that a genetic predisposition or bias towards certain novel object or experience can exists, such situations would be akin to the approach or avoidance-biased configuration in our study. Per the value-based decision-making framework, there is no provision for the selection or avoidance of such actions purely based on salience. However, an action with no approach tendency can be approached due to transient elevation of mDA with no implication to long-term learning. This represents exploratory decisions taken by organisms. Furthermore, at the opposite end, we notice that lower dopamine levels indicate avoidance and the lack of willingness to try a novel action. It is noteworthy that the baseline level of dopamine can be different for people with neurological conditions like depression, Parkinson’s disease, schizophrenia etc.

#### Avoidance-biased configuration

Similarly, in the case of an avoidance-biased paradigm, the action could represent a stimulus the organism has an aversion or fear towards, or has low salience. However, there could be a release of mDA due to a perceived future reward, thereby leading to a transient switch of the avoidance signal to an approach signal. This is indicated by Fig. 5.13, where at baseline (tonic) levels of dopamine, the bias was towards avoidance but with increasing levels of mDA and its modulation of ion channels it becomes possible for the transition of dominance from avoidance to approach. The % DA receptor activation indicated beyond (48,96) shows how strongly an avoidance signal could be switched to an approach signal. Clinically, in individuals under the influence of drugs like cocaine, high dopamine concentrations are present in the striatum. Our data (shown in Fig. 5.13) show how actions, that organisms might normally avoid or be averse to, now become approachable leading to risky and/or unacceptable behaviors. Behavioral accounts of people are also indicative of the fact that under the influence of cocaine they feel motivated and develop delusional self-confidence. Reports in rodents also show that they can be highly motivated to perform tasks they would not be willing to carry out normally (Nguyen et al., 2018; Pascoli et al., 2023).

#### Approach-biased configuration

In case of the approach-biased paradigm, the action request could represent an object of high salience that the organism would like to approach again. However, there could be reduction in extracellular dopamine concentration due to uptake or other mechanisms, that resulting in a transient impact on the decision, causing the organism to avoid this option. This is indicated by Fig. 5.16, where at baseline (tonic) levels of dopamine, the behavioral feature was biased towards approach but with decreasing levels of mDA and its modulation of ion channels it becomes possible for the transition of dominance from approach to avoidance. Furthermore, at very low levels of mDA, even an approach-biased signal switches to a strong avoidance bias signal. This finding may have implications for clinical depression as reports suggest that individuals experience a general lack of motivation to carry out activities in this malaise/disorder. The baseline concentration of dopamine determined by tonic firing of dopaminergic neurons could also vary across different ensembles in the striatum and between individuals, thereby explaining the varying preferences, levels of willingness, or motivational levels of various individuals.

#### Δfrequency (difference in spiking frequency of dMSN and iMSN)

Negative Δfrequency (ΔF) values indicate greater activity of iMSN whereas positive ΔF values indicate greater activity of dMSN. Although in the bias configurations a switch from negative to positive ΔF or vice versa was observed, the key aspect to note is that the synaptic bias was unchanged as the % DA receptor activation varied. Although this is not conclusive evidence of action outcome switch, it is a crucial result that shows how dominance of spiking activity can shift despite the synaptic bias. For more conclusive evidence on action outcome switch, further tests on the effect of this changing spiking activity on the basal ganglia network are required, as attempted in the later part of this study.

### Potential role of cholinergic interneurons (CINs) in regulating response of motivational dopamine (mDA)

We hypothesize that the outcome switch during action selection is potentially driven by the local regulation of dopamine release by cholinergic interneurons (CINs). Glutamatergic inputs from the cortex and thalamus are said to activate these CINs. Let us consider an example where to obtain a future reward, the subject needs to perform few avoidance-biased tasks. A plausible mechanism that is causing the switching of the avoidance-biased tasks as temporarily approachable is the glutamatergic inputs that encode information pertaining to the future reward that trigger the CINs. Experimentally such observations were made where subjects need to exert efforts or go through pain to attain future reward (Salamone et al., 2007), however it has not been used specifically to test the sub-second timescale fluctuations of DA that would result in switch in excitability of dMSN v/s iMSN as shown by our computational study. Future experiments could consider these bias configurations in conjunction with optogenetic manipulation of CINs to trigger the local release of DA during an avoidance-biased action to see if it is approached.

### What dMSN-iMSN pairs could represent and shifting scales from single neurons to ensemble activity

It has been found that if there is similarity between two actions then there is an overlap between the MSN ensembles encoding for these actions (Klaus et al., 2017; Markowitz and Datta, 2020). Furthermore, there is evidence that dMSN and iMSN activity can be segregated into sub-second behavioral motifs or syllables (Markowitz et al., 2018). Some investigators report even that single body parts movement activity in rats is processed by individual MSN in the striatum (Roesch et al., 2009; Coffey et al., 2016). Collating all these lines of evidence, it is highly plausible that an action is comprised of a few distinguishable or abstract behavioral features (or motifs/syllables), each of whose biases are encoded by a dMSN and iMSN acting in pairs by receiving and processing similar information, as was the case in our study.

### Indirect effect of transient dopamine on SNr neurons via striatal ensemble projections

In light of optogenetic evidence that dMSN and iMSN can inhibit and enhance the spiking activity of SNr neurons followed by evidence of convergence of inputs from direct and indirect pathway onto a single SNr neuron, we have projected ensemble activity onto single SNr neurons (Freeze et al., 2013). The ensemble was comprised of 10 such dMSN-iMSN pairs as that is the typical size of the cluster (Barbera et al., 2016). For each trial, a randomized selection of 10 out of 20 dMSN and 10 out of 20 iMSN was made to project its spiking activity onto SNr neurons. In Fig. 4, it was shown how avoidance outcome was attained in an approach-biased neural ensemble at low dopamine whereas approach outcome was attained in an avoidance-biased neural ensemble at elevated dopamine.

The typical SNr response triggered via striatal ensembles when exposed to low dopamine, irrespective of bias, was a lack of significant pause in activity as predicted and is indicative of avoidance outcome. This has strong implications in explaining how in conditions such as depression where there is abnormally low dopamine we are demotivated to perform any action. On the other hand, as the dopamine levels were elevated around the striatal ensembles, the pause in SNr activity started becoming significant. As in indicated in Fig. 5, when % DA receptor activation was increased for the ensemble neurons of both no-bias and avoidance-bias configurations, we observed a leftward shift in the onset time of pause as well as an increase in the maximum duration of pause. Longer duration of pauses could indicate longer window for the selection of action whereas earlier onset of pause could indicate quicker selection of this action over other competing actions.

In the qualitative heat map (Fig. 6), we observe that at baseline activation, the outcome clearly matches the respective bias of the ensemble. Furthermore, at abnormal elevations of DA i.e. greater than phasic DA release that was reported by (Dreyer et al., 2010) to be ∼(24,96)%, we actually observe how avoidance-biased actions are also being favoured for approach. This could be the case under the influence of drugs such as cocaine that prevents dopamine reuptake, leading to abnormally elevated levels in the striatum. The levels of transient DA could signal the effort needed towards an action that also depends on the bias, greater bias indicates greater deviation from tonic DA is needed to switch the signal from avoidance to approach or vice versa.

### Limitations and future work

Stylized morphology was used in our models to limit morphological variability and focus on the biophysical effects on excitability. In future studies, realistic morphologies could be used with additional focus on structural plasticity changes. Furthermore, in relation to this, the effect of dopamine modulation on spine clusters formations in MSN would be an interesting topic to consider. Another consideration is to include lateral connections between dMSN and iMSN as they could have a direct impact on the competition between these two sub-types. This was accounted for in our models to some extent wherein the GABAergic input connections implicitly represent some lateral connectivity with other MSNs.

Although signal transmission delay was accounted for both the direct and indirect pathway, the model has scope for improvement in including intermediate mechanisms for the indirect pathway. We were limited by the resources at our disposal at this stage to implement this but is definitely an important consideration for future improvements to this computational framework. Furthermore, the biases within the striatal neural ensemble might not be to the same extent for each dMSN-iMSN pairing. If dMSN-iMSN pairs truly encode for behavioural motifs (or syllables) then the bias for each such unit would be different. This means that even if the overall bias towards an action is approach or avoidance, there could be non-uniform biases between dMSN-iMSN pairs. This is a possibility to be aware of but it won’t change how transient dopamine affects the network when in comes to switch in outcome. Experimental testing specifically on how cholinergic interneurons can cause transient motivational responses leading to action outcome switch despite the neural ensemble bias is warranted in future studies.

